# Population Genetics and Analysis of Molecular Variance (AMOVA) of the *Monkeypox* virus interferon-alpha-beta receptor gene and its evolutionary relationship with the *Orthopoxvirus* genus

**DOI:** 10.1101/2022.09.02.506281

**Authors:** Pierre Teodosio Felix, Ana Clara da Silva Santos, Iasmin Auta do Nascimento, Lidiane Santos da Silva

**Author notes:** Corresponding author/ **Contact**.

## Abstract

In this work are used 59 haplotypes of the gene of the interferon-alpha-beta receptor of Monkeypox virus, Buffalopox virus, Camelpox virus, Cowpox virus, Ectromelia virus, Rabbitpox virus, Vaccinia virus and Variola virus, recuperated from GENBANK/NCBI. All sequences were publicly available on the Platform of the National Center for Biotechnology Information (NCBI) and were analyzed for their Molecular Variance (AMOVA), haplotypic diversity, mismatch, demographic and spatial expansion, molecular diversity and time of evolutionary divergence. The results suggested that there was a high diversity among haplotypes, with high numbers of transitions, transversions and mutations of the indels type for 5 of the 8 groups analyzed and with slight population expansion perceived in the neutrality tests. The estimators used in this study did not support a uniformity among all the results found, which ensures the lack of conservation of the gene, as well as its protein product, a fact that stimulates the use of therapies based on neutralizing antibodies and the development of new drugs that act as adjuvants to the function of the interferon-alpha-beta receptor gene.

## 1. Introduction

Monkeypox (MPXV) is an Orthopoxvirus (genus covering camelpox, cowpox, vaccinia, and smallpox virus), and has two strains of existing genomes: one in West Africa (giving rise to a less severe stage of the disease in humans and non-human primates) and another in Central Africa (which selectively silences the transcription of genes involved in immunity of the disease in severe hosts). (Reynolds *et al*, 2006; Hammarlund *et al*, 2008).

Although endemic to these regions, the Monkeypox virus has been spreading to other African countries, having as its first recorded case that of a 9-month-old baby in the Democratic Republic of Congo in 1970 (Breman *et al*, 1980). Come able from the 1970s and 1980s, there was an increase in confirmed cases in 9 times and in 2003 cases have been reported in other countries, however, the Democratic Republic of Congo (DRC) is the most affected in the period of 5 decades (Bunge *et al*, 2022). Nigeria enters as the second most affected country, resulting from the 181 confirmed cases, with an outbreak reported in September 2017, from this outbreak was diagnosed cases in Israel and Singapore, cases that originated in Nigeria through travel to these countries, including the United Kingdom with three confirmed cases as originating in Nigeria (Erez *et al*, 2019). The clinical picture of MPXV is usually similar to that of smallpox, and fever, headache, fatigue, rashes, and in addition, the appearance of lymphadenopathy (enlarged lymph nodes that function as filters for harmful substances) can be observed, presented only in Monkeypox. Thus, it is suggested that the presence of lymphadenopathy is an important indicator of immunological recognition of the presence of the virus. (Damon, 2011).

Over evolutionary time, *Orthopoxviruses* adapted to limit the immune response in the human host. Viral strategies to avoid immunological interference may influence even the adverse effects of smallpox vaccination. This occurred through mechanisms that weakened the action of effectors that had the function, inhibiting antiviral effects, which conferred immunoregulatory activity. Among these, we mention <u>interferons</u> and their cellular receptors. (Stanford *et al*, 2007; Randall and Goodbourn, 2008).

Interferons (IFNs) are natural cell signaling glycoproteins that belong to the cytokine class and that participate in cell control and replication, besides being modifiers of the immune response, with antiviral, antiproliferative and immunomodulatory effects (Priyanka *et al*, 2014). IFNs can be divided into three distinct groups. Type I includes IFN-α (alpha) and β (beta), produced by epithelial cells and fibroblasts, contribute to the first line of antiviral defense (Ank *et al*, 2006) specific receptors of its cell surface, called IFN-α / β receptor (IFNAR), is present in all type I IFNs, and consists of the IFNAR1 and IFNAR2 chains. IFN-ε (epsilon), IFN-κ (kappa) and IFN-ω (omega) also fall under type I and are present in humans (Ank *et al*, 2006).

Human IFN type II includes only IFN-γ (gamma), also called immune IFN. It is induced by porcytokines such as interleukin 12 (IL-12) and produced by auxiliary T lymphocytes (Th1 CD4+) and CD8+ T stimulated by foreign antigens, and also “natural killer” (NK) cells. IFN-γ involved in the regulation of immunological and inflammatory responses and has some antiviral and antitumor effects. One of its main functions is to potentiate the effects of IFN type I by recruiting leukocytes to an infection site and stimulating macrophages to eliminate bacteria (Goodbourn *et al*, 2000). Type III interferons include three members, IFN-λ (lambda) 1, 2 and 3, also known as IL-29, IL-28A and IL-28B, respectively. All types of IFNs have the ability to increase the expression of proteins of the main histocompatibility complex (MHC) class I and, thus, to promote CD8+ T-cell responses (Teixeira *et al*, 2008).

Poxviruses encode proteins that serve as invasion strategies in the immunological functions of interferon (IFN), and one of them is the expression of the IFN/BP protein that prevents the interaction of IFN with cell receptors. Detailed interaction studies were performed with VARV and MPXV, and the expression of IFN/BP to identify the adaptation to the human IFN system, where it was possible to observe that the Monkeypox strain efficiently inhibited the antiviral activity of IFN-i types tested, as well as the smallpox strain also specifically inhibited the IFN-i activity. (Smith and Alcami, 2002; Alcami *et al*, 2000).

Progress in molecular biology and genomics has improved understanding of viral infection and monkeypox replication, which has been shown to be a virus with a relatively large genome, with about 196,858 base pairs and which form a good part of the genetic material needed for viral replication in the cellular cytoplasm. Viral types and strains differ in the entry of the virus that occurs immediately after an interaction between several viral ligands and receptors on the cell surface, for example, condroitin sulfate and heparano sulfate (Chung *et al*, 1998; Lin *et al*, 2000). When in the cytoplasm, the virus releases prepackaged proteins that deactivate defenses and encourage gene expression early (Munyon *et al*, 1967; Fonseca *et al*, 2003). Poxviruses are considered self-sufficient in relation to other viruses, but have a very limited range of hosts, which proposes dependence on host elements (Werden *et al*, 2008; McFadde *et al*, 2005). Microarrays have been used in genome recognition and profiling with a special focus on understanding the dynamics of viral gene expression and pathogenicity. However, few studies have used tools to test the host response to infections with poxvirus, in general, specifically in the case of Monkeypox. The microarrays allow the study of alterations caused by MPV in genic expression of epithelial cells of a *kidney of the mulatta monkey* identifying the main connections of interaction between virus and host, this infection caused modulation about 2,702 genes of the hosts. In some data analyses, the regulated genes are grouped into different functional groups, thus canonical pathways and networks that can be linked to viral biogenesis (Abdulnaser *et al*, 2010).

With regard to vaccines, studies indicate that smallpox vaccination provides protection by Orthopoxvirus, indicating that individuals who received routine vaccination, even those who took it before eradication in 1980, may still be protected against Monkeypox infection, as well as those who have not yet received the vaccine may be contributing to the spread of the disease (Symons *et al*, 1995).

Nevertheless, a better understanding of the factors involved in the transmission and dissemination of Monkeypox is still necessary, such as the importance and need for prevention information and care that should be taken in public health. As well as greater effort in demonstrating the protein affinities of VARV and MPXV and their immunoregulatory potentials, such as, in evolutionary time, the adaptation of viruses to a human host. (Babkin and Babkina, 2022).

Trying to understand evolutionary aspects of the Interferon gene and its probable behavior in Monkeypox infection, we from the Laboratory of Population Genetics and Computational Evolutionary Biology (LaBECom-UNIVISA)perform a Molecular Variance Analysis in a PopSet with 59 sequences of the interferon-alpha-beta receptor gene, available in the database of the National Biotechnology Information Center (NCBI).

## 2. Objectives

Test the existing molecular variance levels in 59 haplotypes of the interferon-alpha-beta receptor gene, departing *from Monkeypox virus, Buffalopox virus, Camelpox virus, Cowpox virus, Ectromelia virus, Rabbitpox virus, Vaccinia virus* and *Variola virus*.

## 3. Methodology

### Databank

59 haplotypes of the interferon-alpha-beta gene of *Monkeypox virus, Buffalopox virus, Camelpox virus, Cowpox virus, Ectromelia virus, Rabbitpox virus, Vaccinia virus* and *Variola virus*, were recovered from GENBANK (https://www.ncbi.nlm.nih.gov/popset/60547302?report=fasta) on August 6, 2022. These haplotypes had 1429 base pairs, but once aligned using the MEGA X program (KUMAR *et al*., 2018), had ambiguous sites, lost data and gaps excluded, as well as all sites conserved, resulting in a 100% polymorphic sequence with 213 bp of extension, which served as material for analysis.

### 3.1 To visualize the variable sites

The graphical representation of the sites was made using the WEBLOGO v3 software, described by CROOKS *et al*., 2004.

### 3.2 Genetic structural analyses

Paired F_ST_ estimators, Molecular Variance (AMOVA), Genetic Distance, incompatibility, demographic and spatial expansion analyses, molecular diversity and evolutionary divergence time were obtained with the Arlequin V. 3.5 Software (EXCOFFIER *et al*., 2010) using 1000 random permutations (KUMAR *et al*, 2018). The geographic distance and F_ST_ matrices were not compared. All the steps in this process are described below:

#### Genetic diversity

Among the routines of LaBECom, this test is used to measure the genetic diversity equivalent to the heterozygosity expected in the groups studied. We used for this the standard index of genetic diversity H, described by Nei (1987). This can also be estimated by the method proposed by PONS and PETIT (1995).

#### Site Frequency Spectrum (SFS)

According to LaBECom protocols, we used this local frequency spectrum analytical test (SFS), from DNA sequence data that allow us to estimate the demographic parameters of the frequency spectrum. Simulations are made using fastsimcoal2 software, available in http://cmpg.unibe.ch/software/fastsimcoal2/.

#### Molecular diversity indices

Molecular diversity indices are obtained by means of the average number of paired differences, as described by Tajima in 1993, in this test we use sequences that do not fit the neutral theory model that establishes the existence of a balance between mutation and genetic drift.

#### Calculating THETA estimators

Population parameters are used in our Laboratory when we want to qualify the genetic diversity of the populations studied. These estimates, classified as Theta Hom – which aim to estimate the expected homozygosity in a population in balance between drift and mutation and the estimates Theta (S) (WATTERSON, 1975), Theta (K) (EWENS, 1972) and Theta (π) (TAJIMA, 1983).

#### Calculation of the incompatibility distribution

In LaBECom, analyses of the distribution of incompatibility are always performed relating the observed number of differences between pairs of haplotypes, trying to define or establish a pattern of population demographic behavior, as already described by (ROGERS; HARPENDING, 1992; HUDSON and SLATKIN, 1991; RAY *et al*., 2003, EXCOFFIER, 2004).

#### Pure demographic expansion

This model is always used when we intend to estimate the probability of differences observed between two haplotypes not recombined and randomly chosen, this methodology in our laboratory is used when we assume that the expansion, in a haploid population, reached a momentary balance even having passed through generations of τ, from sizes 0 N to 1 N. In this case, the probability of observing the S differences between two haplotypes not recombined and randomly chosen is given by the probability of observing two haplotypes with S differences in this population (Watterson, 1975).

#### Space expansion

The use of this model in LaBECom is generally indicated if the reach of a population is initially restricted to a very small area, and when one notices signs of growth of the same, in the same space and in a relatively short time. The resulting population usually becomes subdivided in the sense that individuals tend to mate with geographically close individuals rather than random individuals. To follow the dimensions of spatial expansion, we at LaBECom always take into account:

L: Number of loci

Gamma correction: This fix is always used when mutation rates do not seem uniform for all locations.

nd: Number of substitutions observed between two DNA sequences.

ns: Number of transitions observed between two DNA sequences.

nv: Number of transversions observed between two DNA sequences.

ω: G + C ratio, calculated in all DNA sequences of a given sample.

Paired difference: Shows the number of loci for which two haplotypes are different.

Percentage difference: This difference is responsible for producing the loci percentage for which two haplotypes are different.

#### Haplotypic inferences

We use these inferences for haplotypic or genotypic data with unknown gametic phase. Following our protocol, inferences are estimated by observing the relationship between haplotype i and xi times its number of copies, generating an estimated frequency (^pi). With genetic data with unknown gametic phase, haplotype frequencies are estimated by the maximum probability method, and can also be estimated using the expected Maximization (DM) algorithm.

#### Jukes and Singer Method

This method, when used in LaBECom, allows estimating a corrected percentage of how different two haplotypes are. This correction allows us to assume that there have been several substitutions by site, since the most recent ancestor of the two haplotypes studied. Here, we also assume a correction for identical replacement rates for all four nucleotides A C, G and T.

#### Kimura method with two parameters

Like the previous test, this fix allows for multiple site overrides, but takes into account different override rates between transitions and transversions.

#### Method Tamura (1992)

We at LaBECom understand this method as an extension of the 2-parameter Kimura method, which also allows the estimation of frequencies for different haplotypes. However, transient-transversion relationships as well as general nucleotide frequencies are calculated from the original data.

#### Tajima and Nei Method

At this stage, we were also able to produce a corrected percentage of nucleotides for which two haplotypes are different, but this correction is an extension of the Jukes and Cantor method, with the difference of being able to do this from the original data.

#### Model Tamura and Nei

As in the kimura models 2 parameters the distance of Tajima and Nei, this correction allows, inferring different rates of transversions and transitions, besides being able to distinguish transition rates between purines and pyrimidines.

#### Network of minimum coverage

To calculate the distance between the OTU (operational taxonomic units) of the paired distance matrix of the haplotypes, a minimum spanning network (MSN) tree was used, with a slight modification of the algorithm described in Rohlf (1973). We usually use free software written in Pascal called MINSPNET. EXE running in dos language, previously available at: http://anthropologie.unige.ch/LGB/software/win/min-span-net/.

#### Genotypic data with unknown gametic phase

##### EM algorithm

To estimate haplotypic frequencies, we used the maximum probability model with an algorithm that maximizes the expected values. The use of this algorithm in LaBECom allows to obtain the maximum probability estimates from multilocal data of the gametic phase (phenotypic data). It is a slightly more complex procedure, as it does not allow us to make a simple genetic count, since individuals in a population can be heterozygous to more than one locus.

##### ELB algorithm

Very similar to the previous algorithm, ELB attempts to reconstruct the gametic (unknown) phase of multilocal genotypes by adjusting the sizes and locations of neighboring loci to explore some rare recombination.

#### Neutrality tests

##### Ewens-Watterson homozygosis test

We use this test in LaBECom for haploid and diploid data. This test is used only as a way to summarize the distribution of the allelic frequency, without taking into account its biological significance. This test is based on the sample theory of all neutral Ewens alleles (1972) and tested by Watterson (1978). It is now limited to sample sizes of 2,000 genes or less and 1,000 different alleles (haplotypes) or less. It is still used to test the hypothesis of selective neutrality and population equilibrium against natural selection or the presence of some advantageous all links.

##### Accurate Ewens-Watterson-Slatkin Test

This test created by Slatikin in 1991 and adapted by himself in 1996. is used in our protocols when we want to compare the probabilities of random samples with those of observed samples.

##### Population Amalgam Chakraborty Test

This test was proposed by Chakrabordy in 1990, serves to calculate the observed probability of a neutral sample randomly with a number of all links equal to or greater than that observed, based on the infinite alleles model and the sample theory of Neutral All links of Ewens (1972).

##### Tajima Selective Neutrality Test

We use this test in our Laboratory when DNA sequences or haplotypes produced by RFLP are short. It is based on the 1989 Tajima test, using the model of infinite locations without recombination. It travels two estimators using the mutation as a parameter.

##### FS FU Selective Neutrality Test

Also based on the model of infinite sites without recombination, the FU test is suitable for short DNA sequences or haplotypes produced by RFLP. However, in this case, it evaluates the observed probability of a randomly neutral sample with a number of alleles equal to or less than the observed value. In this case, the estimator used is φ.

#### Methods that measure interpopulation diversity

##### Genetic structure of the population inferred by molecular variance analysis (AMOVA)

This stage is the most used in the LaBECom protocols because it allows to know the genetic structure of populations that measure their variances, this methodology, first defined by Cockerham in 1969 and 1973) and, later adapted by other researchers, is essentially similar to other approaches based on genetic frequency variance analyses, but takes into account the number of mutations among haplotypes. When the population group is defined, we can define a specific genetic structure that will be tested, that is, we can create a hierarchical analysis of variance by dividing the total variance into covariance components, being able to measure intra-individual differences, interindividual differences and/or interpopulated differences.

##### Minimum Coverage Network (MSN) among haplotypes

In LaBECom, this tree is generated using operational taxonomic units (OTU). This tree is calculated from the paired distance matrix using a modification of the algorithm described in Rohlf (1973).

##### Locus-per-locus AMOVA

We performed this analysis for each locus separately, as it is performed at the haplotypic level and the components of variance and f statistics are estimated for each locus that generates separately in a more global panorama.

##### Paired genetic distances between populations

This is the most present analysis in LaBECom’s work. These generate paired FST parameters that are always used, extremely reliable, to estimate the short-term genetic distances between the populations studied, in this model a slight algorithmic adaptation is applied to linearize the genetic distance with the time of population divergence **(Reynolds *et al*. 1983**; Slatkin, 1995).

##### Reynolds Distance (Reynolds *et al*. 1983)

Here we measured how much fixed pairs of n-size haplotypes diverged over t generations, based on F_ST_ indices.

##### Slatkin Linearized FST’s (Slatkin 1995)

We used this test in LaBECom when we want to know how much two N-size haploid populations diverged for generations behind a population of identical size and managed to remain isolated and without migration. This is a demographic model and applies very well to the phylogeography work of our Laboratory.

##### Average number of differences between populations

In this test, we assumed that the relationship between the gross (D) and liquid (AD) number of differences between populations is the increase in genetic distance between populations (Nei and Li, 1979).

##### Relative population sizes: divergence between populations of unequal sizes

We use this method in LaBECom when we want to estimate the time of divergence between populations of equal sizes (Gaggiotti and Excoffier, 2000), assuming that two populations diverged from an ancestral population of N0 size a few generations, and that they have remained isolated from each other ever since. In this method we assume that, although the sizes of the two infant populations are different, the sum of them will always correspond to the size of the ancestral population. The procedure is based on the comparison of intra- and inter-population (π) diversities that have great variance, which means that, due to short divergence times, the average diversity found in the population may be higher than that observed among populations. These calculations should therefore be made if the assumptions of a pure fission model are met and if the divergence time is relatively old. The results of this simulation show that this procedure leads to better results than other methods that do not take into account unequal population sizes, especially when the relative sizes of the daughter populations are in fact unequal.

##### Accurate population differentiation tests

We at LaBECom understand that this test is an analog of Fisher’s exact test in a 2×2 contingency table extended to a rxk contingency table. It was described in Raymond and Rousset (1995) and tests the hypothesis of a random distribution of k different haplotypes or genotypes among r populations.

##### Assignment of individual genotypes to populations

Inspired by what had been described in Paetkau *et al* (1995, 1997) and Waser and Strobeck (1998) this method determines the origin of specific individuals, knowing a list of potential populations of origin and uses the allelic frequencies estimated in each sample of its original constitution.

##### Loci detection under selection by F statistics

We use this test when we suspect that natural selection affects genetic diversity among populations. This method was adapted by Cavalli-Sforza in 1996 from a 1973 work by Lewontin and Krakauer.

## 4. Results

### 4.1. General properties of interferon-alpha-beta receptor gene sequences in the orthopoxviruses studied

59 haplotypes of the interferon-alpha-beta receptor gene of Monkeypox virus, Buffalopox virus, Camelpox virus, Cowpox virus, Ectromelia virus, Rabbitpox virus, Vaccinia virus and Variola virus, were recovered from GENBANK (https://www.ncbi.nlm.nih.gov/popset/60547302) on August 6, 2022 (Table 1).

**Table 1.** Description of all sequences analyzed in this work. By GENBANK for update, access number, and organism.

|  | Update | Definition | Accession | Organism |
| --- | --- | --- | --- | --- |
| 1 | Mar 12, 2005 | Buffalopox virus strain BP-1 soluble interferon-alpha/beta gene receptor, complete cds | AY315835.1 | *Buffalopox virus* |
| 2 | Mar 12, 2005 | Camelpox virus strain CP Saudi soluble interferon-alpha/beta receptor gene, complete cds | AY315878.1 | *Camelpox Virus* |
| 3 | Mar 12, 2005 | Camelpox virus strain CP-1 soluble interferon-alpha/beta gene receptor, complete cds | AY315874.1 | *Camelpox Virus* |
| 4 | Mar 12, 2005 | Camelpox virus strain CP-1231 soluble interferon-alpha/beta receptor gene, complete cds | AY315875.1 | *Camelpox Virus* |
| 5 | Mar 12, 2005 | Camelpox virus strain CP-1260 soluble interferon-alpha/beta gene receptor, complete cds | AY315876.1 | *Camelpox Virus* |
| 6 | Mar 12, 2005 | Camelpox virus strain CP-5 soluble interferon-alpha/beta gene receptor, complete cds | AY315877.1 | *Camelpox Virus* |
| 7 | Mar 12, 2005 | Cowpox virus strain Brighton soluble interferon-alpha/beta gene receptor, complete cds | AY315836.1 | *Cowpox Virus* |
| 8 | Mar 12, 2005 | Cowpox virus strain EP-1 soluble interferon-alpha/beta receptor gene, complete cds | AY315843.1 | *Cowpox Virus* |
| 9 | Mar 12, 2005 | Cowpox virus strain EP-2 soluble interferon-alpha/beta gene receptor, complete cds | AY315844.1 | *Cowpox Virus* |
| 10 | Mar 12, 2005 | Cowpox virus strain EP-267 soluble interferon-alpha/beta gene receptor, complete cds | AY315845.1 | *Cowpox Virus* |
| 11 | Mar 12, 2005 | Cowpox virus strain EP-3 soluble interferon-alpha/beta gene receptor, complete cds | AY315846.1 | *Cowpox Virus* |
| 12 | Mar 12, 2005 | Cowpox virus strain EP-4 soluble interferon-alpha/beta gene receptor, complete cds | AY315847.1 | *Cowpox Virus* |
| 13 | Mar 12, 2005 | Cowpox virus strain EP-5 soluble interferon-alpha/beta gene receptor, complete cds | AY315848.1 | *Cowpox Virus* |
| 14 | Mar 12, 2005 | Cowpox virus strain EP-6 soluble interferon-alpha/beta gene receptor, complete cds | AY315849.1 | *Cowpox Virus* |
| 15 | Mar 12, 2005 | Cowpox virus strain EP-7 soluble interferon-alpha/beta gene receptor, complete cds | AY315850.1 | *Cowpox Virus* |
| 16 | Mar 12, 2005 | Cowpox virus strain EP-8 soluble interferon-alpha/beta gene receptor, complete cds | AY315851.1 | *Cowpox Virus* |
| 17 | Mar 12, 2005 | Cowpox virus strain EXP-GR soluble interferon-alpha/beta gene receptor, complete cds | AY315852.1 | *Cowpox Virus* |
| 18 | Mar 12, 2005 | Cowpox virus strain Gruzia soluble interferon-alpha/beta gene receptor, complete cds | AY315837.1 | *Cowpox Virus* |
| 19 | Mar 12, 2005 | Cowpox virus strain Ham-85 soluble interferon-alpha/beta gene receptor, complete cds | AY315838.1 | *Cowpox Virus* |
| 20 | Mar 12, 2005 | Cowpox virus strain OPV-88 soluble interferon-alpha/beta gene receptor, complete cds | AY315842.1 | *Cowpox Virus* |
| 21 | Mar 12, 2005 | Cowpox virus strain Puma M-73 soluble interferon-alpha/beta gene receptor, complete cds | AY315839.1 | *Cowpox Virus* |
| 22 | Mar 12, 2005 | Cowpox virus strain Rev-91 soluble interferon-alpha/beta receptor gene, complete cds | AY315840.1 | *Cowpox Virus* |
| 23 | Mar 12, 2005 | Cowpox virus strain RP-1 soluble interferon-alpha/beta gene receptor, complete cds | AY315853.1 | *Cowpox Virus* |
| 24 | Mar 12, 2005 | Cowpox virus strain RP-9 soluble interferon-alpha/beta gene receptor, complete cds | AY315854.1 | *Cowpox Virus* |
| 25 | Mar 12, 2005 | Cowpox virus strain Turk-74 soluble interferon-alpha/beta gene receptor, complete cds | AY315841.1 | *Cowpox Virus* |
| 26 | Mar 12, 2005 | Ectromelia virus strain K1 clone 2 soluble interferon-alpha/beta receptor gene, complete cds | AY315885.1 | *Ectromelia virus* |
| 27 | Mar 12, 2005 | Ectromelia virus strain K1 soluble interferon-alpha/beta gene receptor, complete cds | AY315884.1 | *Ectromelia virus* |
| 28 | Mar 12, 2005 | Monkeypox virus strain CDC# v70-I-187 soluble interferon-alpha/beta receptor gene, complete cds | AY315882.1 | *Monkeypox Virus* |
| 29 | Mar 12, 2005 | Monkeypox virus strain CDC# v78-I-3945 soluble interferon-alpha/beta receptor gene, complete cds | AY315883.1 | *Monkeypox Virus* |
| 30 | Mar 12, 2005 | Monkeypox virus strain CDC# v79-I-005 soluble interferon-alpha/beta receptor gene, complete cds | AY315880.1 | *Monkeypox Virus* |
| 31 | Mar 12, 2005 | Monkeypox virus strain CDC# v97-I-004 soluble interferon-alpha/beta receptor gene, complete cds | AY315881.1 | Monkeypox Virus |
| 32 | Mar 12, 2005 | Monkeypox virus strain Congo 8 soluble interferon-alpha/beta gene receptor, complete cds | AY315879.1 | Monkeypox Virus |
| 33 | Mar 12, 2005 | Rabbitpox virus strain RPU soluble interferon-alpha/beta receptor gene, complete cds | AY315834.1 | Rabbitpox virus |
| 34 | Mar 12, 2005 | Vaccinia virus strain Chambon St-Yves Menard soluble interferon-alpha/beta receptor gene, complete cds | AY315830.1 | Vaccinia virus |
| 35 | Mar 12, 2005 | Vaccinia virus strain Copenhagen soluble interferon-alpha/beta receptor gene, complete cds | AY315828.1 | Vaccinia virus |
| 36 | Mar 12, 2005 | Vaccinia virus strain Copenhagen soluble interferon-alpha/beta receptor gene, complete cds | AY315828.1 | Vaccinia virus |
| 37 | Mar 12, 2005 | Vaccinia virus strain CVI-78 soluble interferon-alpha/beta receptor gene, complete cds | AY315833.1 | Vaccinia virus |
| 38 | Mar 12, 2005 | Vaccinia virus strain EM-63 soluble interferon-alpha/beta receptor gene, complete cds | AY315832.1 | Vaccinia virus |
| 39 | Mar 12, 2005 | Vaccinia virus strain LIVP soluble interferon-alpha/beta gene receptor, complete cds | AY315829.1 | Vaccinia virus |
| 40 | Mar 12, 2005 | Vaccinia virus strain Patwadanger soluble interferon-alpha/beta receptor gene, complete cds | AY315831.1 | Vaccinia virus |
| 41 | Mar 12, 2005 | Variola virus strain 12/62 soluble interferon-alpha/beta receptor gene, complete cds | AY315860.1 | Variola virus |
| 42 | Mar 12, 2005 | Variola virus strain 22/62 soluble interferon-alpha/beta receptor gene, complete cds | AY315855.1 | Variola virus |
| 43 | Mar 12, 2005 | Variola virus strain 6/58 soluble interferon-alpha/beta gene receptor, complete cds | AY315856.1 | Variola virus |
| 44 | Mar 12, 2005 | Variola virus strain Aslam soluble interferon-alpha/beta receptor gene, complete cds | AY315861.1 | Variola virus |
| 45 | Mar 12, 2005 | Variola virus strain Aziz soluble interferon-alpha/beta receptor gene, complete cds | AY315862.1 | Variola virus |
| 46 | Mar 12, 2005 | Variola virus strain Brazil 128 soluble interferon-alpha/beta receptor gene, complete cds | AY315857.1 | Variola virus |
| 47 | Mar 12, 2005 | Variola virus strain Brazil 131 soluble interferon-alpha/beta receptor gene, complete cds | AY315858.1 | Variola virus |
| 48 | Mar 12, 2005 | Variola virus strain Butler soluble interferon-alpha/beta receptor gene, complete cds | AY315863.1 | Variola virus |
| 49 | Mar 12, 2005 | Variola virus strain Congo-2 soluble interferon-alpha/beta receptor gene, complete cds | AY315864.1 | Variola virus |
| 50 | Mar 12, 2005 | Variola virus strain Congo-9 soluble interferon-alpha/beta receptor gene, complete cds | AY315865.1 | Variola virus |
| 51 | Mar 12, 2005 | Variola virus strain Ind-3a soluble interferon-alpha/beta receptor gene, complete cds | AY315866.1 | Variola virus |
| 52 | Mar 12, 2005 | Variola virus strain India 164 soluble interferon-alpha/beta receptor gene, complete cds | AY315859.1 | Variola virus |
| 53 | Mar 12, 2005 | Variola virus strain India 378 soluble interferon-alpha/beta gene receptor, complete cds | AY315867.1 | Variola virus |
| 54 | Mar 12, 2005 | Variola virus strain Kuw-5 soluble interferon-alpha/beta gene receptor, complete cds | AY315868.1 | Variola virus |
| 55 | Mar 12, 2005 | Variola virus strain M-Abr-60 soluble interferon-alpha/beta gene receptor, complete cds | AY315869.1 | Variola virus |
| 56 | Mar 12, 2005 | Variola virus strain M-Sok-60 soluble interferon-alpha/beta receptor gene, complete cds | AY315870.1 | Variola virus |
| 57 | Mar 12, 2005 | Variola virus strain M-Sur-60 soluble interferon-alpha/beta receptor gene, complete cds | AY315871.1 | Variola virus |
| 58 | Mar 12, 2005 | Variola virus strain Ngami soluble interferon-alpha/beta receptor gene, complete cds | AY315872.1 | Variola virus |
| 59 | Mar 12, 2005 | Variola virus strain Semat soluble interferon-alpha/beta receptor gene, complete cds | AY315873.1 | Variola virus |

These haplotypes had 1429 base pairs, but once aligned using the MEGA X program (KUMAR *et al*., 2018), had their ambiguous sitios, lost data, gaps and all preserved sites excluded, resulting in a 100% polymorphic sequence with 213bp extension. The graphic representation of these sites could be seen in a logo built with the software WEBLOGO 3. (CROOKS *et al*., 2004), where the size of each nucleotide is proportional to its frequency for certain sites. (Figure 1).

**Figure 1:**
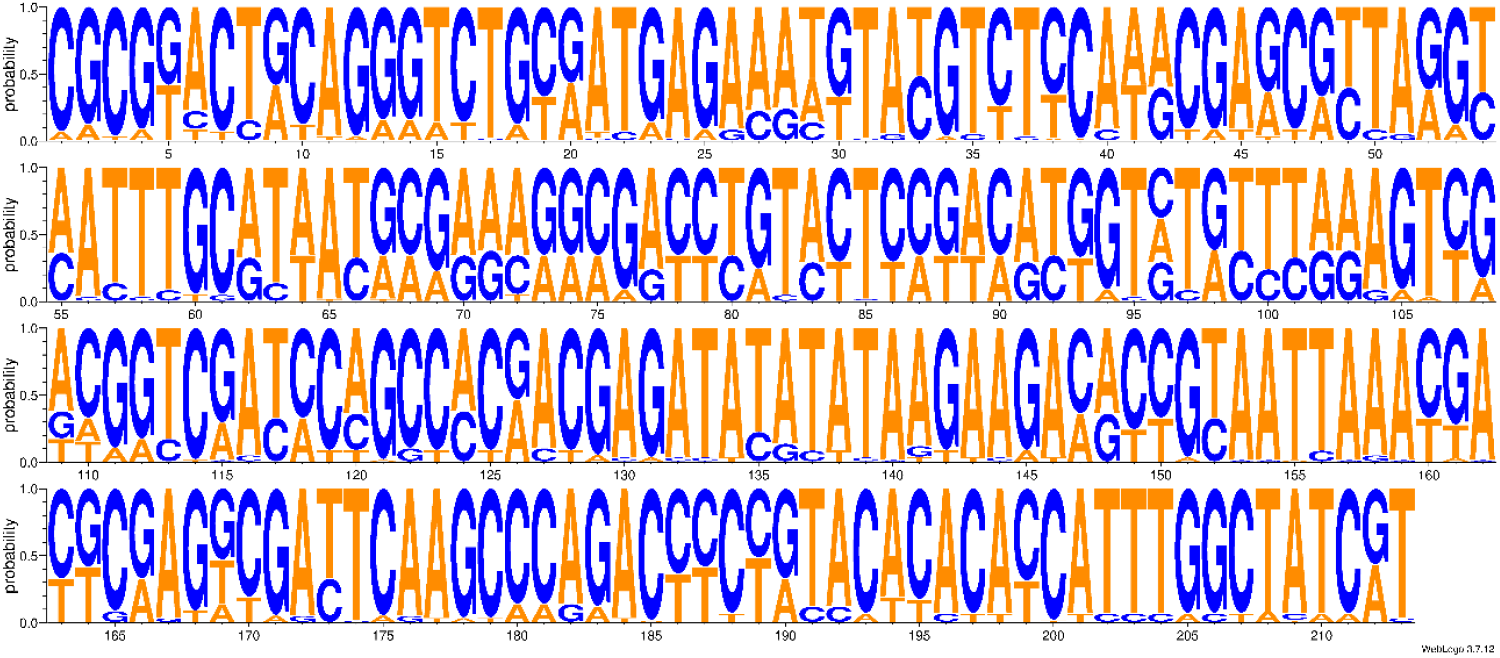
Graphic representation of 213 polymorphically sites of interferon-alpha-beta receptor gene sequences.

### 4.2. Molecular Variance Analysis (AMOVA) and Genetic Distance

Genetic distance and molecular variation **(AMOVA)** analyses were highly significant for the groups studied, presenting a variation component of 25% between populations and 8% within populations. The F_ST_ value (76%) showed a high fixation index, with highly significant evolutionary divergences within and between groups (Table 2 and Figures 2 and 3). There were also no significant similarities for the time of genetic evolutionary divergence among all populations; supported by τ variations, mismatch analyses and demographic and spatial expansion analyses (Tables 3,4 and 5); (Figures 4, 5 and 6).

**Tabela 2.** Components of haplotypic variation and paired FST value for the 59 haplotypes of the interferon-alpha-beta receptor gene.

| Source of variation | d.f. | Sum of squares | Variance components | Percentage of variation |
| --- | --- | --- | --- | --- |
| Among populations | 7 | 1174.747 | 25.44290 Va | 76.03 |
| Within populations | 50 | 400.977 | 8.01954 Vb | 23.97 |
| Total | 57 | 1575.724 | 33.46244 |  |
| Fixation Index | **$F_{ST}$:** | **0.76034** |  |  |

Significance tests (1023 permutations)

**Figure 2.**
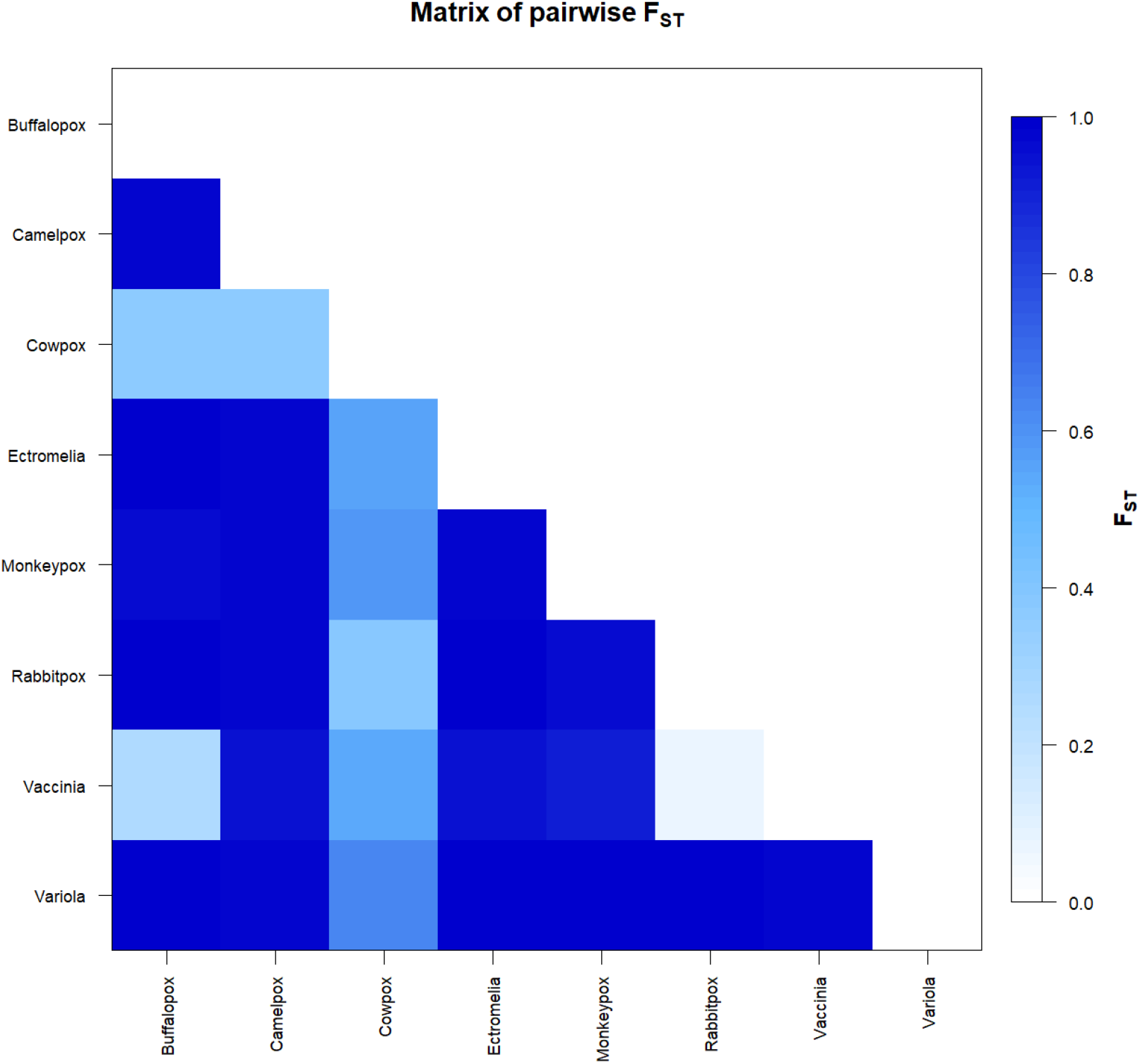
Genetic distance matrix based on F_ST_ among the 8 groups of Orthopoxvirus studied. * Generated by the statistical package in R language using the output data of the Software Arlequin version 3.5.1.2.

**Figure 3.**
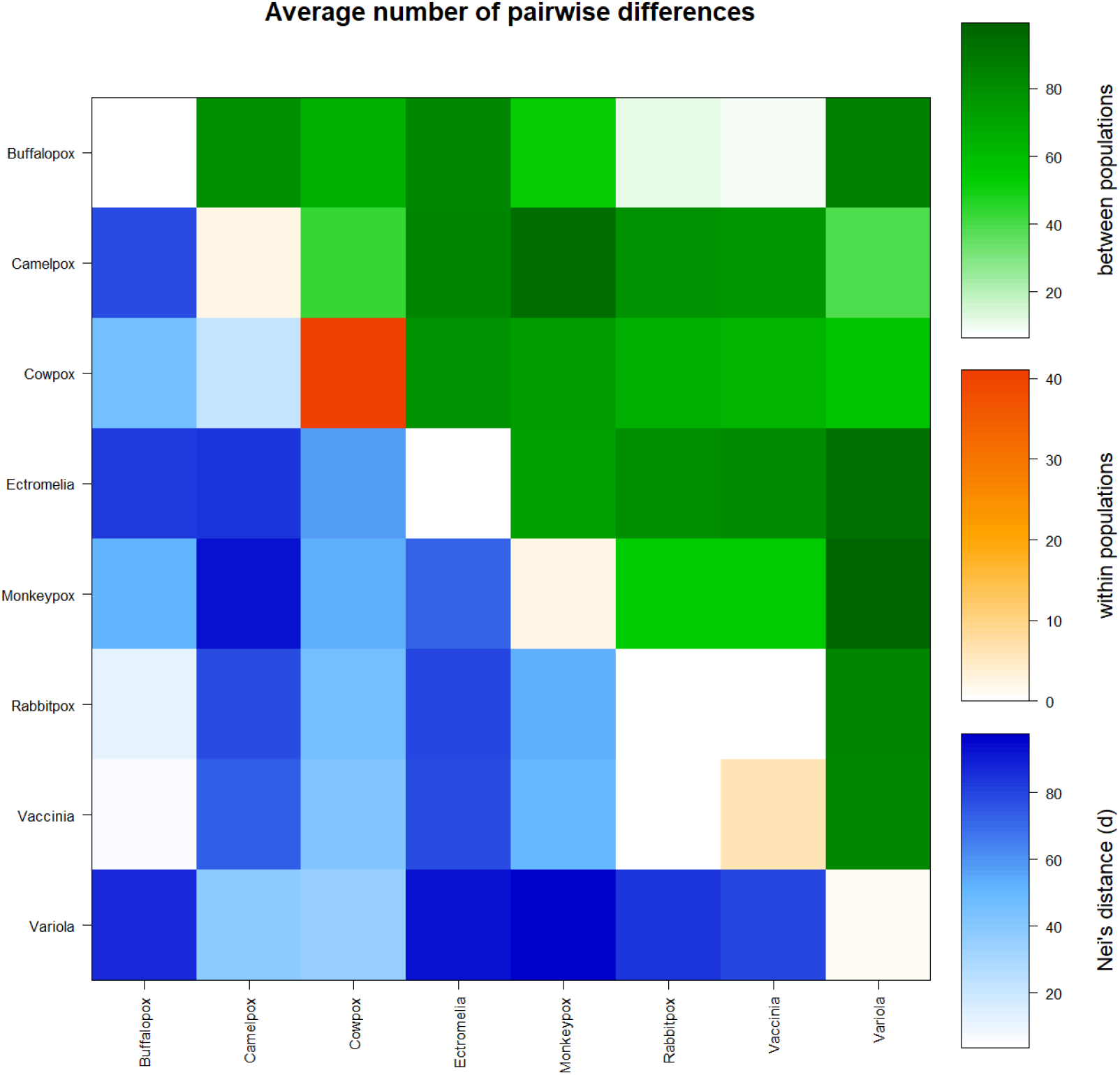
Matrix of paired differences between the populations studied: between the groups; within the groups; and Nei distance for the 8 groups of Orthopoxvirus studied. * Generated by the statistical package in R language using the output data of the Software Arlequin version 3.5.1.2.

**Tabela 3.** Molecular Diversity Indices for the 8 Groups of Orthopoxvirus studied

| Statistics | Buffalopox | Camelpox | Cowpox | Ectromelia | Monkeypox | Rabbitpox | Vaccinia | Variola | Mean | s.d. |
| --- | --- | --- | --- | --- | --- | --- | --- | --- | --- | --- |
| No. of transitions | 0 | 3 | 75 | 0 | 3 | 0 | 13 | 3 | 12.125 | 25.760 |
| No. of transversions | 0 | 3 | 45 | 0 | 2 | 0 | 2 | 0 | 6.500 | 15.602 |
| No. of substitutions | 0 | 6 | 120 | 0 | 5 | 0 | 15 | 3 | 18.625 | 41.265 |
| No. of indels | 0 | 0 | 0 | 0 | 0 | 0 | 0 | 0 | 0.000 | 0.000 |
| No. of ts. sites | 0 | 3 | 74 | 0 | 3 | 0 | 13 | 3 | 12.000 | 25.411 |
| No. of tv. sites | 0 | 3 | 45 | 0 | 2 | 0 | 2 | 0 | 6.500 | 15.602 |
| No. of subst. sites | 0 | 6 | 115 | 0 | 5 | 0 | 15 | 3 | 18.000 | 39.511 |
| Total: 139 |  |  |  |  |  |  |  |  |  |  |
| No. private subst. sites | 0 | 6 | 111 | 0 | 5 | 0 | 10 | 2 | 16.750 | 38.250 |
| No. of indel sites | 0 | 0 | 0 | 0 | 0 | 0 | 0 | 0 | 0.000 | 0.000 |
| Pi | 0.000 | 2.400 | 41.076 | 0.000 | 2.400 | 0.000 | 6.067 | 0.725 | 6.58348 | 14.08825 |
| Theta_k | N.A. | N.A. | N.A. | N.A. | N.A. | N.A. | N.A. | N.A. | N.A. | N.A. |
| Theta_k_lower | N.A. | N.A. | N.A. | N.A. | N.A. | N.A. | N.A. | N.A. | N.A. | N.A. |
| Theta_k_upper | N.A. | N.A. | N.A. | N.A. | N.A. | N.A. | N.A. | N.A. | N.A. | N.A. |
| Theta_H | N.A. | N.A. | N.A. | N.A. | N.A. | N.A. | N.A. | N.A. | N.A. | N.A. |
| s.d. Theta_H | N.A. | N.A. | N.A. | N.A. | N.A. | N.A. | N.A. | N.A. | N.A. | N.A. |
| Theta_S | 0.00000 | 2.88000 | 32.90313 | 0.00000 | 2.40000 | 0.00000 | 6.56934 | 0.85834 | 5.70135 | 11.21555 |
| s.d. Theta_S | 0.00000 | 1.75772 | 11.53582 | 0.00000 | 1.51331 | 0.00000 | 3.42141 | 0.54971 | 2.34725 | 3.89847 |
| Theta_pi | 0.00000 | 2.40000 | 41.07602 | 0.00000 | 2.40000 | 0.00000 | 6.06667 | 0.72515 | 6.58348 | 14.08825 |
| s.d. Theta_pi | 0.00000 | 1.81989 | 20.85916 | 0.00000 | 1.81989 | 0.00000 | 3.88578 | 0.63201 | 3.62709 | 7.09105 |

**Tabela 4.** Neutrality tests for the sequences for the 8 groups of Orthopoxvirus studied

| Statistics | Buffalopox | Camelpox | Cowpox | Ectromelia | Monkeypox | Rabbitpox | Vaccinia | Variola | Mean | s.d. |
| --- | --- | --- | --- | --- | --- | --- | --- | --- | --- | --- |
| **Ewens-Watterson test** |  |  |  |  |  |  |  |  |  |  |
| Sample size | 1 | 5 | 19 | 2 | 5 | 1 | 6 | 19 | 7.25000 | 7.49762 |
| No. of alleles(unchecked) | 1 | 5 | 19 | 2 | 5 | 1 | 6 | 19 | 7.25000 | 7.49762 |
| Observed F value | N.A. | N.A. | N.A. | N.A. | N.A. | N.A. | N.A. | N.A. | N.A. | N.A. |
| Expected F value | N.A. | N.A. | N.A. | N.A. | N.A. | N.A. | N.A. | N.A. | N.A. | N.A. |
| Watterson test: |  |  |  |  |  |  |  |  |  |  |
| Pr(rand F <= obs F) | N.A. | N.A. | N.A. | N.A. | N.A. | N.A. | N.A. | N.A. | N.A. | N.A. |
| Slatkin's exact test P-value | N.A. | N.A. | N.A. | N.A. | N.A. | N.A. | N.A. | N.A. | N.A. | N.A. |
| **Chakraborty's test** |  |  |  |  |  |  |  |  |  |  |
| Sample size | 1 | 5 | 19 | 2 | 5 | 1 | 6 | 19 | 7.25000 | 7.49762 |
| No. of alleles(unchecked) | 1 | 5 | 19 | 2 | 5 | 1 | 6 | 19 | 7.25000 | 7.49762 |
| Obs. homozygosity | 0.00000 | 0.00000 | 0.00000 | 0.00000 | 0.00000 | 0.00000 | 0.00000 | 0.00000 | 0.00000 | 0.00000 |
| Exp. no. of alleles | 0.00000 | 3.07078 | 15.77575 | 1.00000 | 3.07078 | 0.00000 | 4.43052 | 2.97822 | 3.79076 | 5.10286 |
| P(k or more alleles) | N.A. | N.A. | N.A. | N.A. | N.A. | N.A. | N.A. | N.A. | N.A. | N.A. |
| **Tajima's D test** |  |  |  |  |  |  |  |  |  |  |
| Sample size | 1 | 5 | 19 | 2 | 5 | 1 | 6 | 19 | 7.25000 | 7.49762 |
| S | 0 | 6 | 115 | 0 | 5 | 0 | 15 | 3 | 18.00000 | 39.51130 |
| Pi | 0.00000 | 2.40000 | 41.07602 | 0.00000 | 2.40000 | 0.00000 | 6.06667 | 0.72515 | 6.58348 | 14.08825 |
| Tajima's D | 0.00000 | -1.14554 | 1.03409 | 0.00000 | 0.00000 | 0.00000 | -0.47165 | -0.42196 | -0.12563 | 0.61596 |
| Tajima's D p-value | 1.00000 | 0.20000 | 1.00000 | 1.00000 | 0.60000 | 1.00000 | 0.50000 | 0.30000 | 0.70000 | 0.34226 |
| **Fu's FS test** |  |  |  |  |  |  |  |  |  |  |
| No. of alleles(unchecked) | 1 | 5 | 19 | 2 | 5 | 1 | 6 | 19 | 7.25000 | 7.49762 |
| Theta_pi | 0.00000 | 2.40000 | 41.07602 | 0.00000 | 2.40000 | 0.00000 | 6.06667 | 0.72515 | 6.58348 | 14.08825 |
| Exp. no. of alleles | 0.00000 | 3.07078 | 15.77575 | 1.00000 | 3.07078 | 0.00000 | 4.43052 | 2.97822 | 3.79076 | 5.10286 |
| FS | 0.00000 | -2.67988 | -3.62299 | 0.00000 | -2.67988 | 0.00000 | -1.79286 | -340282 | 0.00000 | 0.00000 |
| FS p-value | N.A. | 0.00000 | 0.00000 | 1.00000 | 0.00000 | N.A. | 0.30000 | 0.00000 | N.A. | N.A. |

**Tabela 5.** Demographic and spatial expansion simulations based on the τ, φ and M indices for the 8 orthopoxvirus groups studied

| Statistics | Buffalopox | Camelpox | Cowpox | Ectromelia | Monkeypox | Rabbitpox | Vaccinia | Variola | Mean | s.d. |
| --- | --- | --- | --- | --- | --- | --- | --- | --- | --- | --- |
| **DEMOGRAPHIC EXPANSION** |  |  |  |  |  |  |  |  |  |  |
| Tau | 0.00000 | 2.75781 | 91.81217 | 0.00000 | 2.50000 | 0.00000 | 4.00000 | 1.00000 | 12.75875 | 31.97777 |
| Tau qt 2.5% | 0.00000 | 1.32027 | 47.61698 | 0.00000 | 2.69337 | 0.00000 | 3.01563 | 0.97070 | 6.95212 | 16.47403 |
| Tau qt 5% | 0.00000 | 1.32027 | 47.61698 | 0.00000 | 2.69337 | 0.00000 | 3.01563 | 0.97070 | 6.95212 | 16.47403 |
| Tau qt 95% | 0.00000 | 4.35548 | 94.08171 | 0.00000 | 4.39845 | 0.00000 | 11.42003 | 2.23441 | 14.56126 | 32.3570 |
| Tau qt 97.5% | 0.00000 | 4.35548 | 94.08171 | 0.00000 | 4.39845 | 0.00000 | 11.42003 | 2.23441 | 14.56126 | 32.3570 |
| Theta0 | 0.00000 | 0.00000 | 0.00154 | 0.00000 | 0.00000 | 0.00000 | 5.49999 | 0.00000 | 0.68769 | 1.94446 |
| Theta0 qt 2.5% | 0.00000 | 0.00000 | 0.00000 | 0.00000 | 0.00000 | 0.00000 | 0.00000 | 0.00000 | 0.00000 | 0.00000 |
| Theta0 qt 5% | 0.00000 | 0.00000 | 0.00000 | 0.00000 | 0.00000 | 0.00000 | 0.00000 | 0.00000 | 0.00000 | 0.00000 |
| Theta0 qt 95% | 0.00000 | 0.02635 | 0.00352 | 0.00000 | 0.06325 | 0.00000 | 10.13357 | 0.63456 | 1.35766 | 3.55268 |
| Theta0 qt 97.5% | 0.00000 | 0.02635 | 0.00352 | 0.00000 | 0.06325 | 0.00000 | 10.13357 | 0.63456 | 1.35766 | 3.55268 |
| Theta1 | 0.00000 | 30.35161 | 62.40004 | 0.00000 | 3414.97837 | 0.00000 | 3414.97837 | 3407.18514 | 1291.23 | 1756.60 |
| Theta1 qt 2.5% | 0.00000 | 3.71367 | 66.48561 | 0.00000 | 13.93023 | 0.00000 | 14.69197 | 1.65198 | 12.55918 | 22.6302 |
| Theta1 qt 5% | 0.00000 | 3.71367 | 66.48561 | 0.00000 | 13.93023 | 0.00000 | 14.69197 | 1.65198 | 12.55918 | 22.6302 |
| Theta1 qt 95% | 0.00000 | 54.84339 | 302.43130 | 0.00000 | 33.09075 | 0.00000 | 132.64295 | 53.91127 | 72.11496 | 103.1 |
| Theta1 qt 97.5% | 0.00000 | 54.84339 | 302.43130 | 0.00000 | 33.09075 | 0.00000 | 132.64295 | 53.91127 | 72.11496 | 103.1 |
| SSD | 0.00000 | 0.01192 | 0.05354 | 0.00000 | 0.03080 | 0.00000 | 0.07622 | 0.03229 | 0.02560 | 0.02820 |
| Model (SSD) p-value | 0.00000 | 0.50000 | 0.00000 | 0.00000 | 0.40000 | 0.00000 | 0.10000 | 0.00000 | 0.12500 | 0.20529 |
| Raggedness index | 0.00000 | 0.07000 | 0.04610 | 0.00000 | 0.16000 | 0.00000 | 0.15111 | 0.21525 | 0.08031 | 0.08468 |
| Raggedness p-value | 0.00000 | 0.60000 | 0.00000 | 0.00000 | 0.40000 | 0.00000 | 0.70000 | 0.00000 | 0.21250 | 0.30443 |
| **SPATIAL EXPANSION** |  |  |  |  |  |  |  |  |  |  |
| Tau | 0.00000 | 2.66995 | 92.94542 | 0.00000 | 2.66812 | 0.00000 | 8.38491 | 0.90555 | 13.44674 | 32.24395 |
| Tau qt 2.5% | 0.00000 | 1.46333 | 75.69351 | 0.00000 | 1.51081 | 0.00000 | 3.32054 | 0.95286 | 10.36763 | 26.41959 |
| Tau qt 5% | 0.00000 | 1.46333 | 75.69351 | 0.00000 | 1.51081 | 0.00000 | 3.32054 | 0.95286 | 10.36763 | 26.41959 |
| Tau qt 95% | 0.00000 | 4.27828 | 93.04544 | 0.00000 | 3.84742 | 0.00000 | 7.16871 | 1.65339 | 13.74915 | 32.14336 |
| Tau qt 97.5% | 0.00000 | 4.27828 | 93.04544 | 0.00000 | 3.84742 | 0.00000 | 7.16871 | 1.65339 | 13.74915 | 32.14336 |
| Theta | 0.00000 | 0.01557 | 3.04950 | 0.00000 | 0.01000 | 0.00000 | 0.01000 | 0.01000 | 0.38688 | 1.07587 |
| Theta qt 2.5% | 0.00000 | 0.00072 | 0.00072 | 0.00000 | 0.00072 | 0.00000 | 0.00072 | 0.00072 | 0.00045 | 0.00037 |
| Theta qt 5% | 0.00000 | 0.00072 | 0.00072 | 0.00000 | 0.00072 | 0.00000 | 0.00072 | 0.00072 | 0.00045 | 0.00037 |
| Theta qt 95% | 0.00000 | 0.01186 | 3.67829 | 0.00000 | 1.37238 | 0.00000 | 5.58751 | 0.03800 | 1.33600 | 2.14994 |
| Theta qt 97.5% | 0.00000 | 0.01186 | 3.67829 | 0.00000 | 1.37238 | 0.00000 | 5.58751 | 0.03800 | 1.33600 | 2.14994 |
| M | 0.00000 | 39.43639 | 2.75997 | 0.00000 | 4382.1069 | 0.00000 | 4312.08784 | 7297.57753 | 2004.246 | 2901.1 |
| M qt 2.5% | 0.00000 | 12.57996 | 0.41126 | 0.00000 | 1.37034 | 0.00000 | 4.34360 | 0.95743 | 2.45782 | 4.33851 |
| M qt 5% | 0.00000 | 12.57996 | 0.41126 | 0.00000 | 1.37034 | 0.00000 | 4.34360 | 0.95743 | 2.45782 | 4.33851 |
| M qt 95% | 0.00000 | 39.46163 | 6.30118 | 0.00000 | 40.48921 | 0.00000 | 782.82907 | 13.26664 | 110.29347 | 272.2699 |
| M qt 97.5% | 0.00000 | 39.46163 | 6.30118 | 0.00000 | 40.48921 | 0.00000 | 782.82907 | 13.26664 | 110.29347 | 272.2699 |
| SSD | 0.00000 | 0.01186 | 0.03375 | 0.00000 | 0.02957 | 0.00000 | 0.06909 | 0.03092 | 0.02190 | 0.02405 |
| Model (SSD) p-value | 0.00000 | 0.60000 | 0.60000 | 0.00000 | 0.40000 | 0.00000 | 0.10000 | 0.00000 | 0.21250 | 0.27484 |
| Raggedness index | 0.00000 | 0.07000 | 0.04610 | 0.00000 | 0.16000 | 0.00000 | 0.15111 | 0.21525 | 0.08031 | 0.08468 |
| Raggedness p-value | 0.00000 | 0.80000 | 1.00000 | 0.00000 | 0.50000 | 0.00000 | 0.90000 | 0.20000 | 0.42500 | 0.43012 |

**Figure 4.**
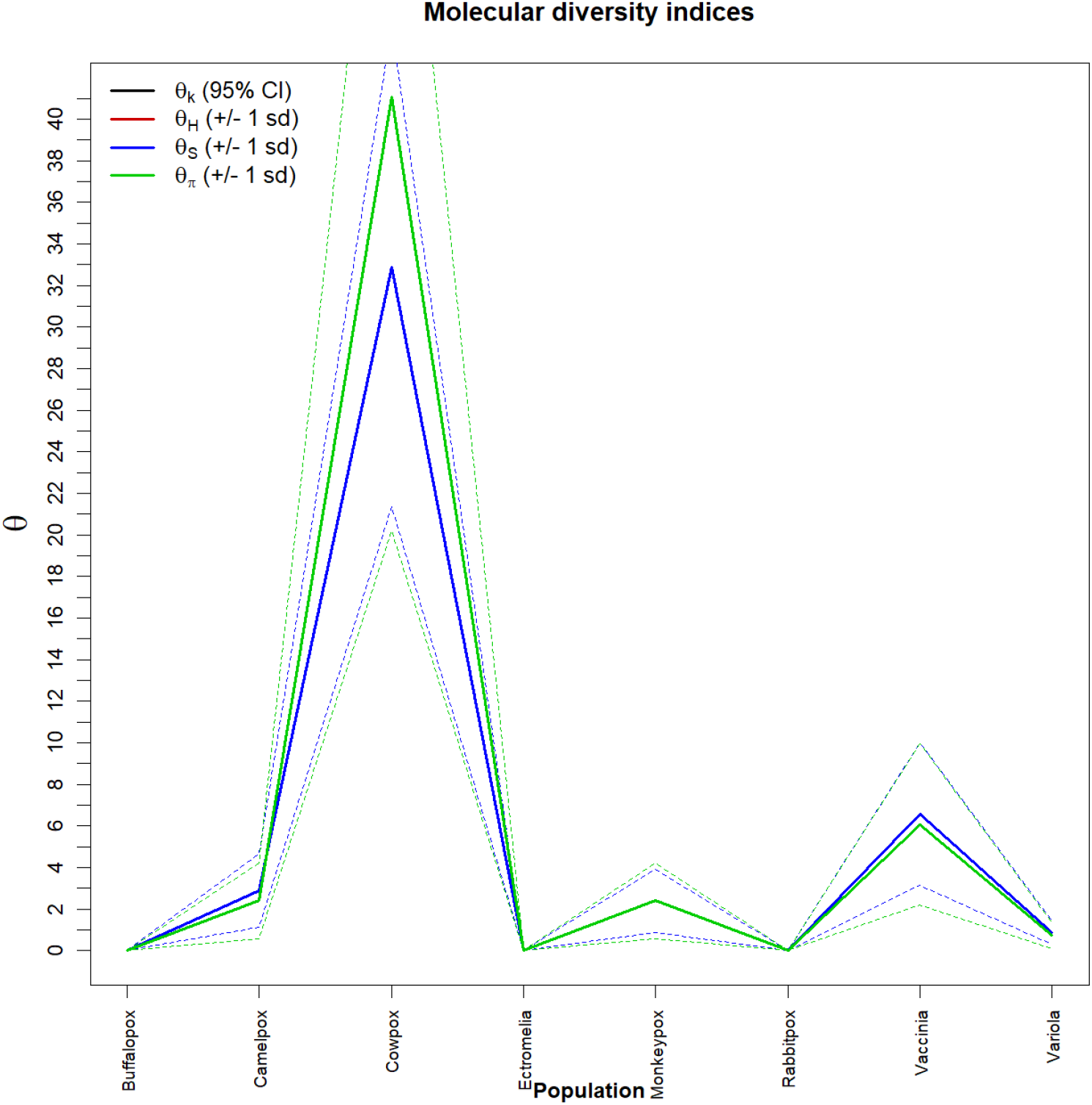
Graph of molecular diversity indices for the 8 groups of Orthopoxvirus studied. In the graph the values φ: (φk) Relationship between the expected number of alleles (k) and the sample size; (φH) Expected homozygosity in a balanced relationship between drift and mutation; (φS) Relationship between the number of segregating sites (S), sample size (n) and non-recombinant sites; (φπ) Relationship between the average number of paired differences (π) and φ. * Generated by the statistical package in Language R using the output data of the Software Arlequin version 3.5.1.2.

**Figure 5.**
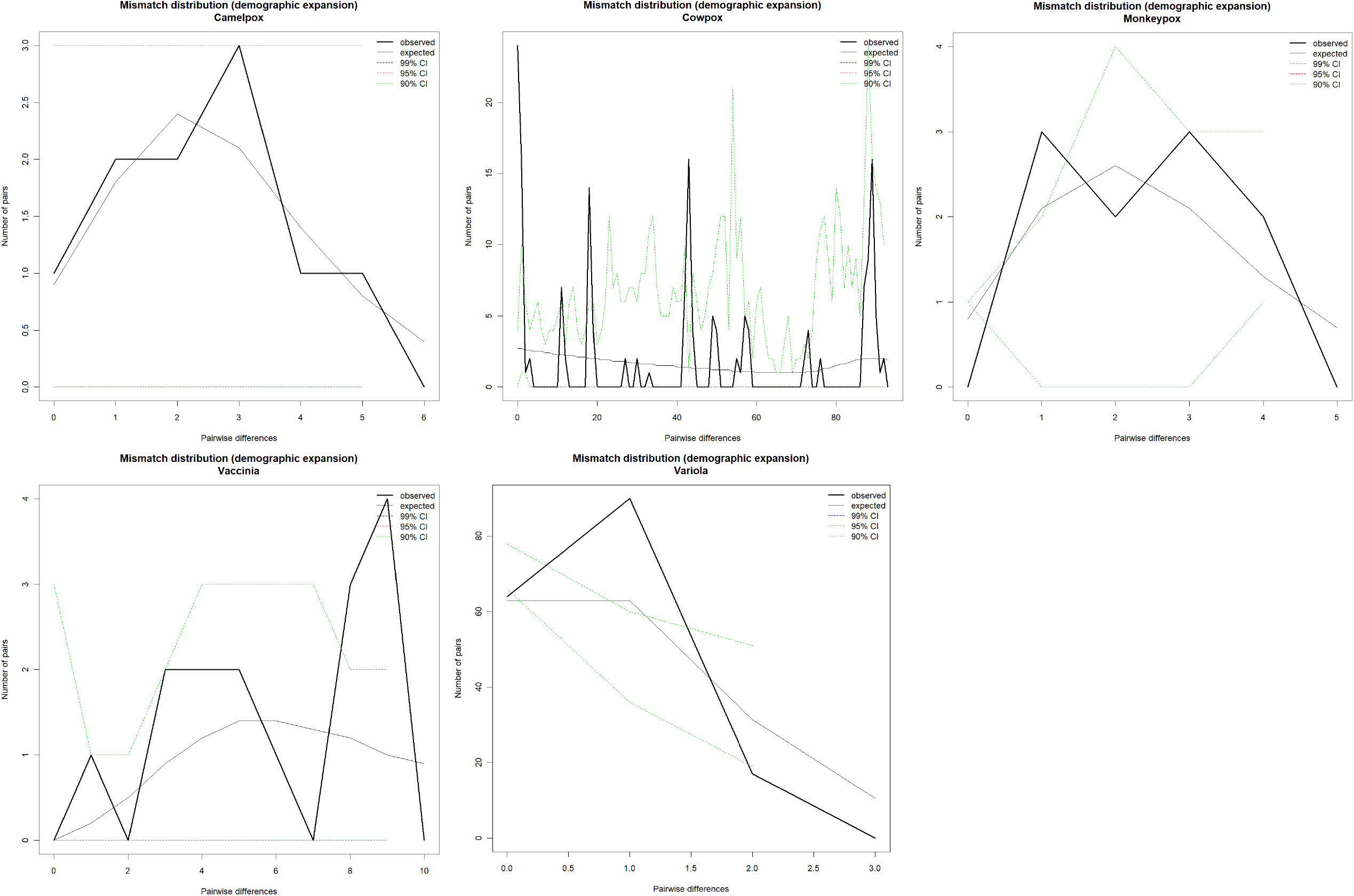
Demographic Expansion Charts for the 5 Groups with The Greatest Molecular Diversity. *Graphs Generated by the R-language statistical package using the output data from version 3.5.1.2 of the Arlequin Software

**Figure 6.**
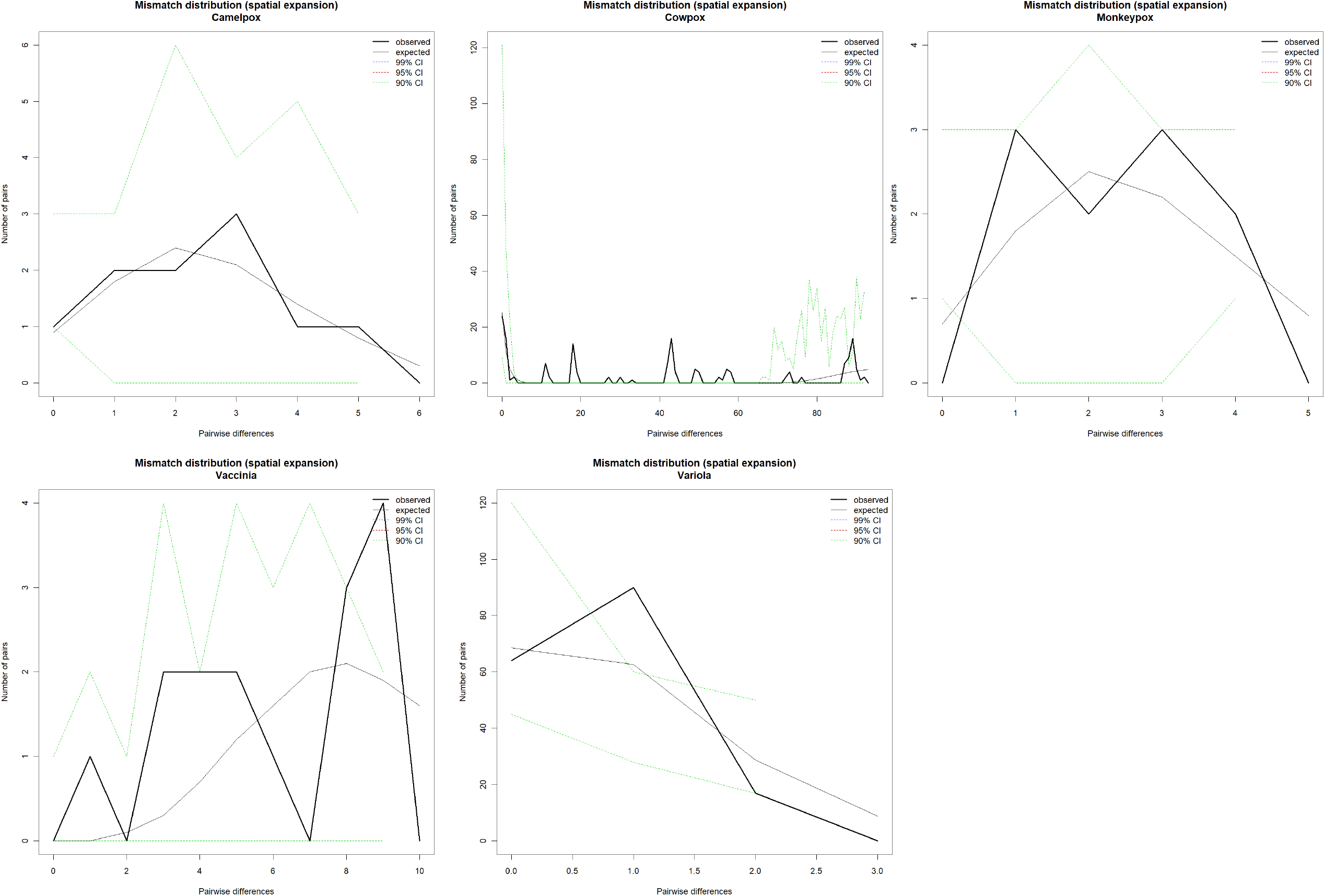
Spatial Expansion Charts for the 5 Groups with The Greatest Molecular Diversity. *Graphs Generated by the R-language statistical package using the output data from version 3.5.1.2 of the Arlequin Software

### 4.3 Analyses of molecular diversity

The molecular analyses estimated by φ reflected a significant level of mutations (transitions and transversions)and 5 m of the 8 groups studied. Indels mutations (insertions or additions) were found in greater numbers only in group Cowpox virus (Table 3); (Figure 4 and 9). The Tajima and FS FU tests showed disagreements between the estimates of general φ and π, but with negative and highly significant values, indicating, once again, the absence of population expansions for all groups except the Cowpox virus group (Table 5) (Figures 5 and 6) . The irregularity index (R= Raggedness) with parametric bootstrap simulated new values φ for before and after a supposed demographic expansion and, in this case, assumed a value equal to zero for the buffalopox virus, Ectromelia virus and Rabbitpox virus groups, probably due to its few molecular diversities (Table 3); (Figure 4). Tau variations (related to the ancestry of all groups) revealed significant divergence s (mainly between the Cowpox virus Camelpox virus and Monkeypox Virus) groups (Figure 7), supported by incompatibility analysis of the observed distribution (τ) (table 5), by the time of divergence and sizes of ancestral populations (Figure 8) and by the inconstant rates of mutation among the eight groups analyzed (Table 4). Nevertheless, a considerable *number of loci*, which are under selection, were detected in our analyses using the F_ST_ matrix (Figure 9).

**Figure 7.**
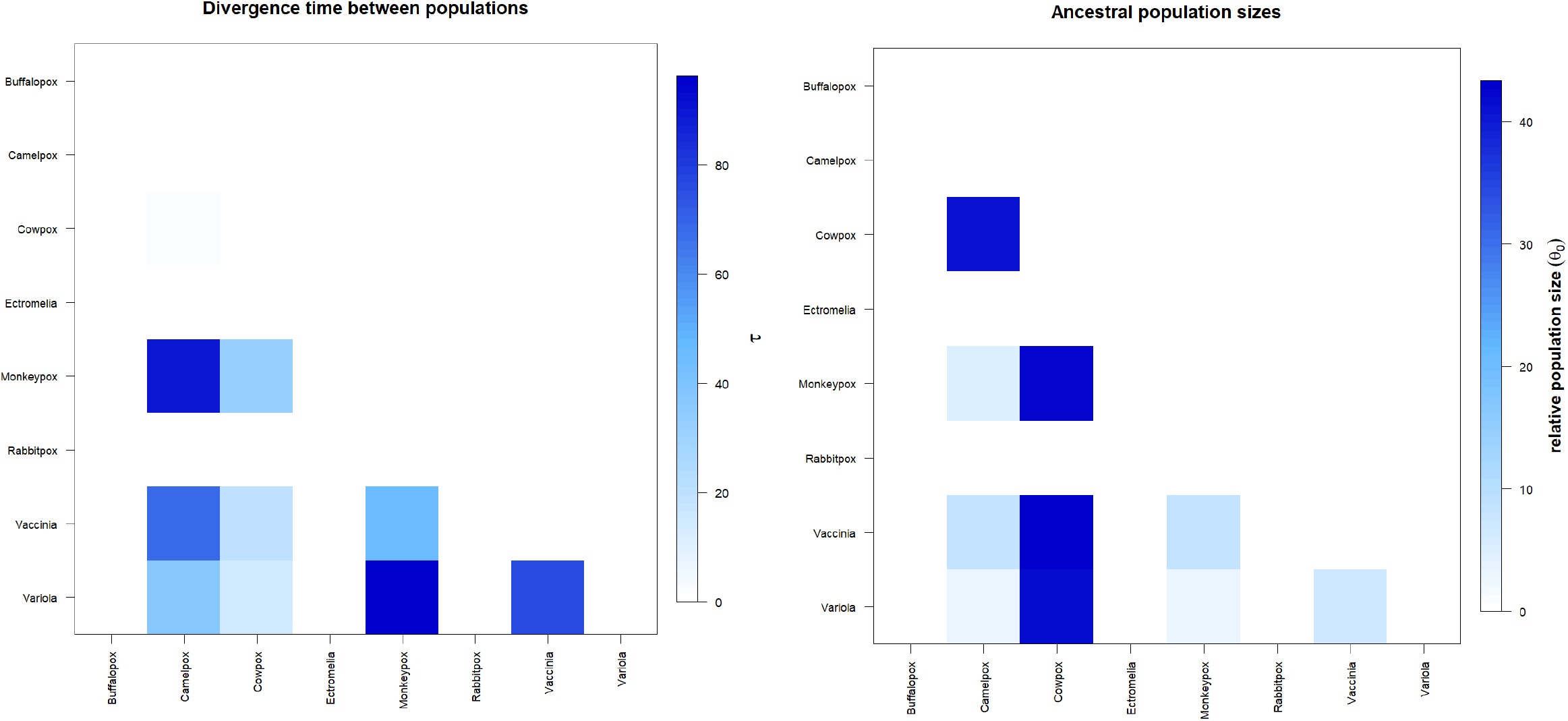
Matrix of divergence time and size of ancestral populations for the 8 groups of Orthopoxvirus studied. In evidence the high value τ present in the monkeypox virus group, Cowpox Virus and Camelpox virus. * Generated by the statistical package in R language using the output data of the Software Arlequin version 3.5.1.2.

**Figure 8.**
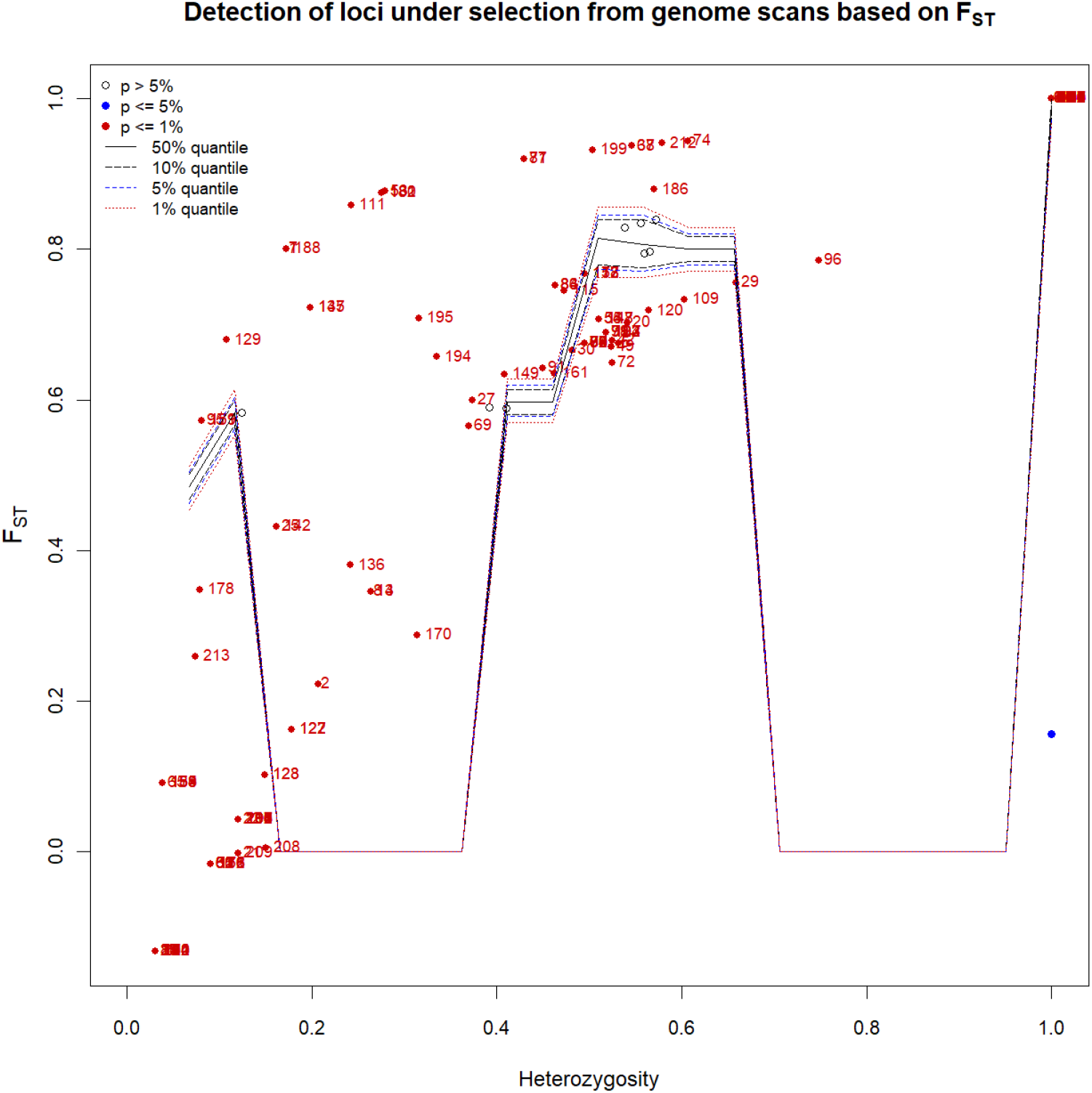
Loci under selection tracked in all genomes using the FST matrix. * Generated by the R-language statistical package using the output data from the Arlequin version 3.5.1.2 software.

**Figure 9.**
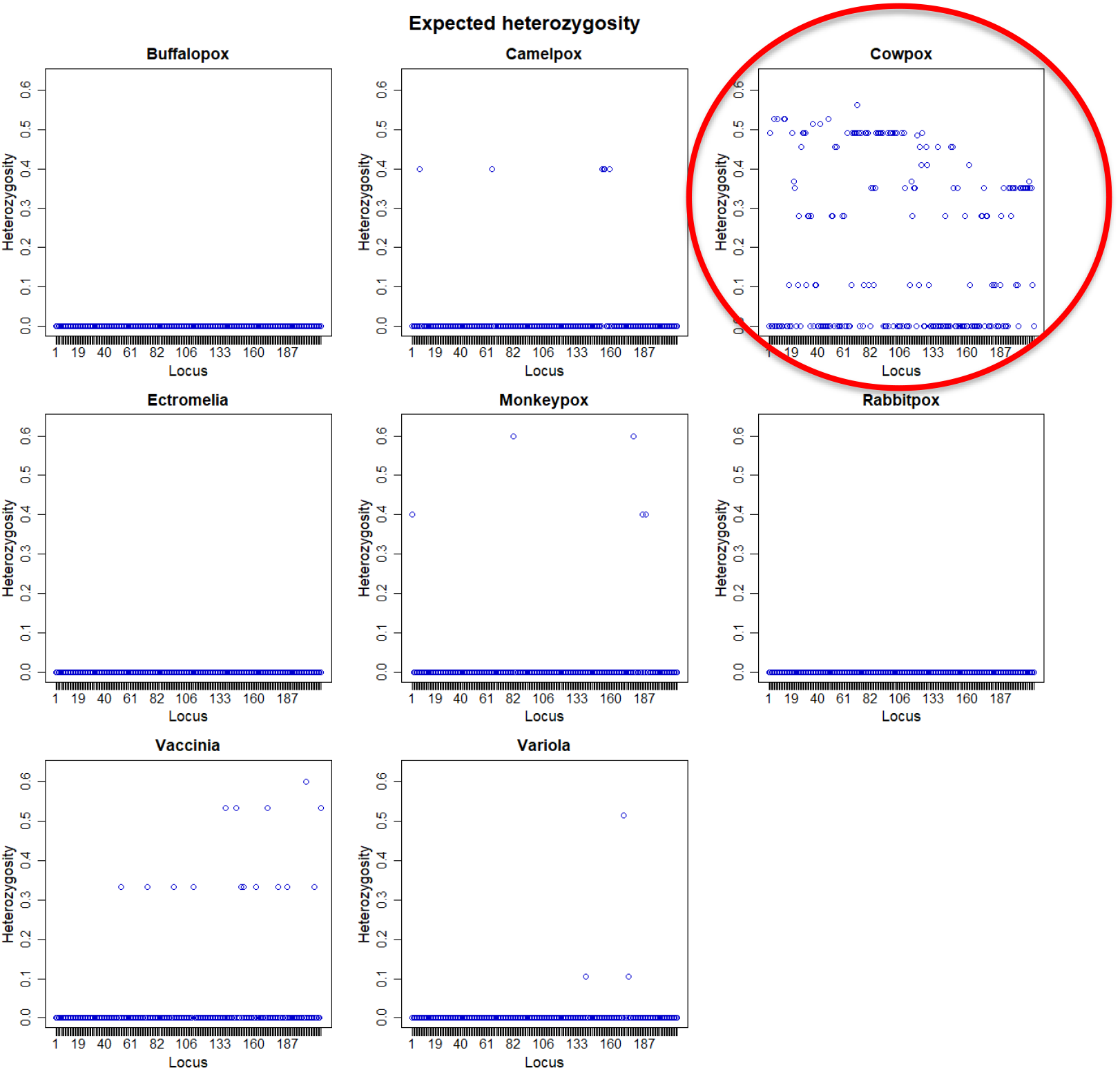
Heterozygosity expected for the 8 groups of Orthopoxvirus studied. In evidence is the high heterozygosity of the Cowpox Virus group. * Generated by the statistical package in R language using the output data of the Software Arlequin version 3.5.1.2.

## 5. Discussion

With the use of molecular variance analysis and molecular population genetics methodologies, which had not yet been used for this PopSet, it was possible to detect the existence of 8 distinct groups of Orthopoxvirus, but with variations among all of them. Different degrees of structuring were detected for each studied group, being essentially higher among one of them (Cowpox). These data suggest that the high degree of structuring present in 5 of the studied groups (Camelpox, Cowpox, Monkeypox, Vaccinia and Variola) may be related to a loss of intermediate haplotypes throughout the generations, possibly associated with an absence of gene flow. These levels of structuring were also supported by methodologies that pointed to a discontinuous pattern of genetic divergence between the groups (discarding the idea of possible geographical sub-isolation stemming from past fragmentation events), taking into account a probable existence of many mutational stages, especially in Cowpox.

The mutations found in the interferon-alpha-beta receptor gene in 5 of the 8 groups studied (Camelpox, Cowpox, Monkeypox, Vaccinia and Variola) seem to have already been fixed by genetic drift probably by the founding effect, which accompanies the behavior of dispersion and/or loss of intermediate haplotypes over generations. The values found for genetic distance support the presence of this continuous pattern of high divergence between the groups studied, since they considered important the minimum differences between the groups when the haplotypes between them were exchanged, as well as the inference of values greater than or equal to that observed in the proportion of these permutations, including the *p value of* the test.

The discrimination of the 59 genetic entities were also perceived by their high inter-haplotypic variations, hierarches in all covariance components: by their intra- and inter-individual differences or by their intra- and intergroup differences, generating a pattern that supports the idea that the significant differences found for the 8 groups, for example, were shared more in their form than in their number, since the result of estimates of the average evolutionary divergence found within these and other countries were very high.

Based on the low level of haplotypic sharing, tests that measure the relationship between genetic distance and geographic distance, such as the Mantel test, should have been performed, but the lack of geographic references of the chosen dataset prevented us from estimating. This lack of correlation leads us to the understanding that the lack of gene flow (observed by non-haplotypic sharing) should be supported firsthand, only by the presence of geographic barriers.

Estimators θ, extremely sensitive to any form of molecular variation (Fu, 1997), did not support the uniformity between the results found by all the methodologies employed, and can be interpreted as a confirmation that there is NO consensus in the conservation of the interferon-alpha-beta receptor gene of Monkeypox virus, Buffalopox virus, Camelpox virus, Cowpox virus, Ectromelia virus, Rabbitpox virus, Vaccinia virus and Variola virus, and it is therefore safe to state that existing polymorphisms, even reflect large variations in their protein products. This consideration provides the safety that the polymorphisms existing in all haplotypes studied bring great differences, further increasing speculation scans about the existence of rapid and silent mutations that exist, as we have shown in this work, significantly increase the genetic variability of all orthopoxvirus studied, making it difficult to work with molecular targets for vaccines and drugs in general. These considerations ensure that the interferon-alpha-beta receptor gene may not be as efficient in suppressing infections and that the significant mutations found may affect the functionality of new drugs that interact as adjuvants in inhibiting or reducing the infectious potential of orthovirus pox.

